# *Drosophila* Male Accessory Gland Displays Developmental Ground Plan of a Primitive Prostate

**DOI:** 10.1101/2021.03.20.436254

**Authors:** Jaya Kumari, Pradip Sinha

**Affiliations:** Department of Biological Sciences and Bioengineering, Indian Institute of Technology Kanpur, Kanpur 208016, India

**Keywords:** Drosophila, Male accessory gland, Prostate, Deep homology

## Abstract

Conservation of developmental genetic toolkits of functionally comparable organs from disparate phyla reveals their deep homology, which may help overcome the challenges of their confounding categorization as either homologous or analogous organs. A male accessory sexual organ in mammals, prostate, for instance, is anatomically disparate from its phylogenetically distant counterpart—the male accessory gland (MAG)—in insects like *Drosophila*. By examining a select set of toolkit gene expression patterns, here we show that *Drosophila* MAG displays deep homology with the mammalian prostate. Like mammalian prostate, MAG morphogenesis is marked by recruitment of fibroblast growth factor receptor, FGFR, a tubulogenesis toolkit signaling pathway, starting early during its adepithelial genesis. Specialization of the individual domains of the developing MAG tube on the other hand is marked by expression of a posterior Hox gene transcription factor, Abd-B, while Hh-Dpp signaling marks its growth. *Drosophila* MAG thus reveals developmental design of unitary bud-derived tube—a ground plan that appears to have been reiteratively co-opted during evolutionary diversification of male accessory sexual organs across distant phylogeny.

## Introduction

Conservation of genetic toolkits—the molecular architects of animal body plan—has helped reveal underlying developmental principles across wide phylogenetic spectrum (Carroll et al., 2001; Shubin et al., 2009). Recruitment of *Pax6* transcription factor for photosensory organs in all bilateria from annelid worms, invertebrates to highly evolved vertebrate eyes being one of the oft-cited and most elegant illustrations of conservation of developmental toolkit genes (Gehring and Seimiya, 2010; Halder et al., 1995; Schwab, 2018). Conservation of developmental toolkit genes—also termed as deep homology (Shubin et al., 1997; Shubin et al., 2009)—has further helped decipher evolutionary relatedness based on genetic blueprints. Deep homology, when revealed, permits deconstructions of complex anatomical features of an organ into its independent or modular components and trace their evolutionary changes (Brandon, 2005; Rasskin-Gutman, 2005). Modularity, rather its breakaways therefore underlie evolutionary diversification from a primitive developmental ground plan of an organ.

The mammalian prostate is derived from a common endodermal embryonic primordium of reproductive and urinary organs termed as the urogenital sinus (UGS) (Marker et al., 2003; Shapiro et al., 2004). Specification of prostatic UGS at its pre-bud stage is followed by its budding, bud elongation, branching and, finally, canalization within each of its primary and secondary branches that create their individual lumens (for review, see Cunha et al., 2018). Epithelial linings of these lumens differentiate into two secretory cell types: luminal and basal (for review, see Marker et al., 2003). Cell fate specification of the prostatic primordium is marked by gain of homeodomain transcription factors, such as Nkx3.1 (for review, see Prins and Putz, 2008), while recruitment of fibroblast growth factor (FGF) and sonic hedgehog (Shh) signalings regulate their growth and branching (Doles et al., 2006; Donjacour et al., 2003; Le et al., 2020; Lin et al., 2007). Posterior Hox genes: for instance, Hoxb13 induces cell differentiation in the luminal cell (Economides and Capecchi, 2003; Huang et al., 2007).

Prostate-like protein-rich seminal fluid secreting organs termed male accessory glands (MAG) are also present in invertebrates, as in the dipteran insect, the fruit fly, *Drosophila* (Findlay et al., 2008; Gilany et al., 2015; Ravi Ram and Wolfner, 2007; Verze et al., 2016). By contrast to endodermal mammalian prostate, *Drosophila* MAG is mesodermal in origin and represents a paired tubular organ (Ahmad and Baker, 2002). It originates from a common adepithelial primordium—which develops in close apposition with the ectodermal male genital disc epithelium—and gives rise to both MAG and seminal vesicle (SV). *Drosophila* counterpart of mammalian FGFR, namely, Breathless (Btl) marks the common embryonic MAG-SV primordium (Ahmad and Baker, 2002). Adult MAG is a relatively simple tubular organ, its lumen is formed by squamous epithelium marked by two differentiated cell types: main and secondary, and is overlaid by circular rings of contractile muscles (Bairati, 1968; Susic-Jung et al., 2012).

Mammalian prostate is proposed to have appeared nearly 65 million years ago (Coffey, 2001) and its evolutionary origin appears to be independent of that of its *Drosophila* counterpart, MAG. Here we show that MAG development in *Drosophila* reveals an essential unitary prostatic tubule and is formed by shared genetic toolkits, which are also recruited during mammalian prostate development. *Drosophila* MAG therefore reveals a primitive and modular developmental ground plan, which appears to have been co-opted across distant phyla.

## Results and Discussion

### Spatial coordinates of developing MAG-SV and genital primordia

In third instar male larvae, a group of mesodermal adepithelial cells—marked by expression of Breathless (Btl)/FGF receptor (FGFR)—migrate and lodge onto the epithelium of the genital imaginal disc and serve as a common primordium of seminal vesicle (SV) and male accessory gland (MAG) (Ahmad and Baker, 2002) (see Figure 1A). The MAG-SV primordium thus develops in close apposition with the genital disc from third larval instar till puparium formation (Figure 1A-C). Optical cross-section further revealed that MAG-SV primordium is surrounded by the genital disc epithelium (Figure 1A’-C’, zoomed cross-section in B’’), the latter marked by a *Drosophila* posterior Hox gene *Abd-B* (Figure1C’, zoomed cross-section in C’’) (Estrada and Sánchez-Herrero, 2001; de Navas et al., 2006) and express its genetic toolkits like *engrailed* (*en*) and *cubitus interruptus* (*ci*) (Figure 1D), which mark its presumptive anterior-posterior compartments wherein *patched (ptc)* (Figure 1E) straddles their boundary while signaling pathways like Wingless (Wg) (Figure 1F) and Decapentapelagic (Dpp) (Figure 1G) regulate its morphogenesis (Casares et al., 1997; Chen and Baker, 1997; Freeland and Kuhn, 1996). The mesodermal MAG-SV and the ectodermal genital primordia thus develop in close apposition prefiguring their mutual contacts in the adult.

**Figure 1.**
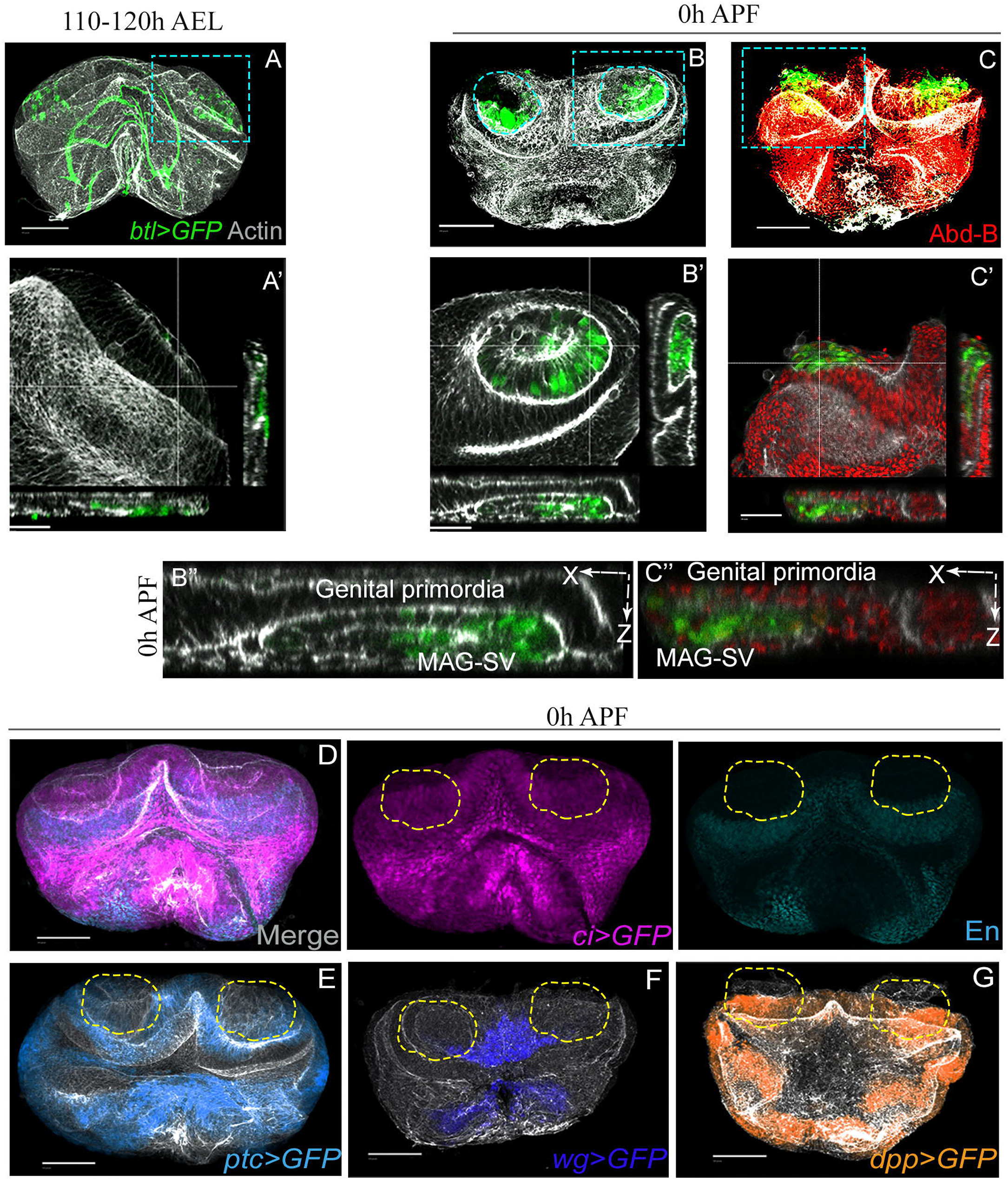
MAG-SV and genital primordia develop in close apposition. (A-C) GFP marked MAG-SV primordium in the backdrop of genital imaginal disc at (A) 3^rd^ instar and (B-C) 0h APF. Zoomed box A’ (Actin, grey) depicts remodelling of the imaginal disc at the 3^rd^ instar, while *btl-Gal4>UAS-GFP* marked MAG-SV primordial cells lodge. Zoomed boxes (B’) show the clustered MAG-SV primordium, (B’’) show the apposition of genital and MAG-SV primordia in X-Z section, and (C’, C’’) the characteristic large nuclei of MAG-SV primordial cells. X-Z view at 0h APF in B’ and C’ is further enlarged below.(D-G) Genital disc pattering genes displayed by their psuedo-colored GFP and immunostaining (En) depict the spatial position of overlaying MAG-SV primordium, which is denoted by broken lines. Scale bar: 100μm Abbreviations: AEL=After egg laying, APF=After puparium formation

### Spatio-temporal expressions of Btl/FGFR and Abd-B during the course of MAG development

Mammalian (rodent) prostate develops from multiple buds that spatially form its bilaterally symmetrical anterior, dorso-lateral, and ventral lobes (Lin et al., 2003; Marker et al., 2003; Timms, 2008). FGFR expression during prostate development is seen selectively during growth and branching morphogenesis; its expression being robust at the distal tips of these branching buds revealing its critical role as a toolkit signaling pathway during rodent prostate development (Huang et al., 2005), Thus, upon conditional knockdown of FGFR, the anterior and the ventral lobes are selectively lost while ductal patterning of dorso-lateral lobe compromised (Lin et al., 2007). In all prostatic lobes, the posterior Hox, Hoxb13 is specifically expressed in the ductal epithelium, its highest expression being the luminal cells (Economides and Capecchi, 2003; Huang et al., 2007). MAG-SV primordium at late 3^rd^ instar, too, was shown to express the Btl/FGFR (Ahmad and Baker, 2002) toolkit signaling pathway while the sole *Drosophila* posterior Hox gene, *Abd-B* (Coiffier et al., 2008) is selectively expressed in the differentiated secondary cells of the adult MAG (Gligorov et al., 2013) (48h after eclosion (AE), Figure 2A).

**Figure 2.**
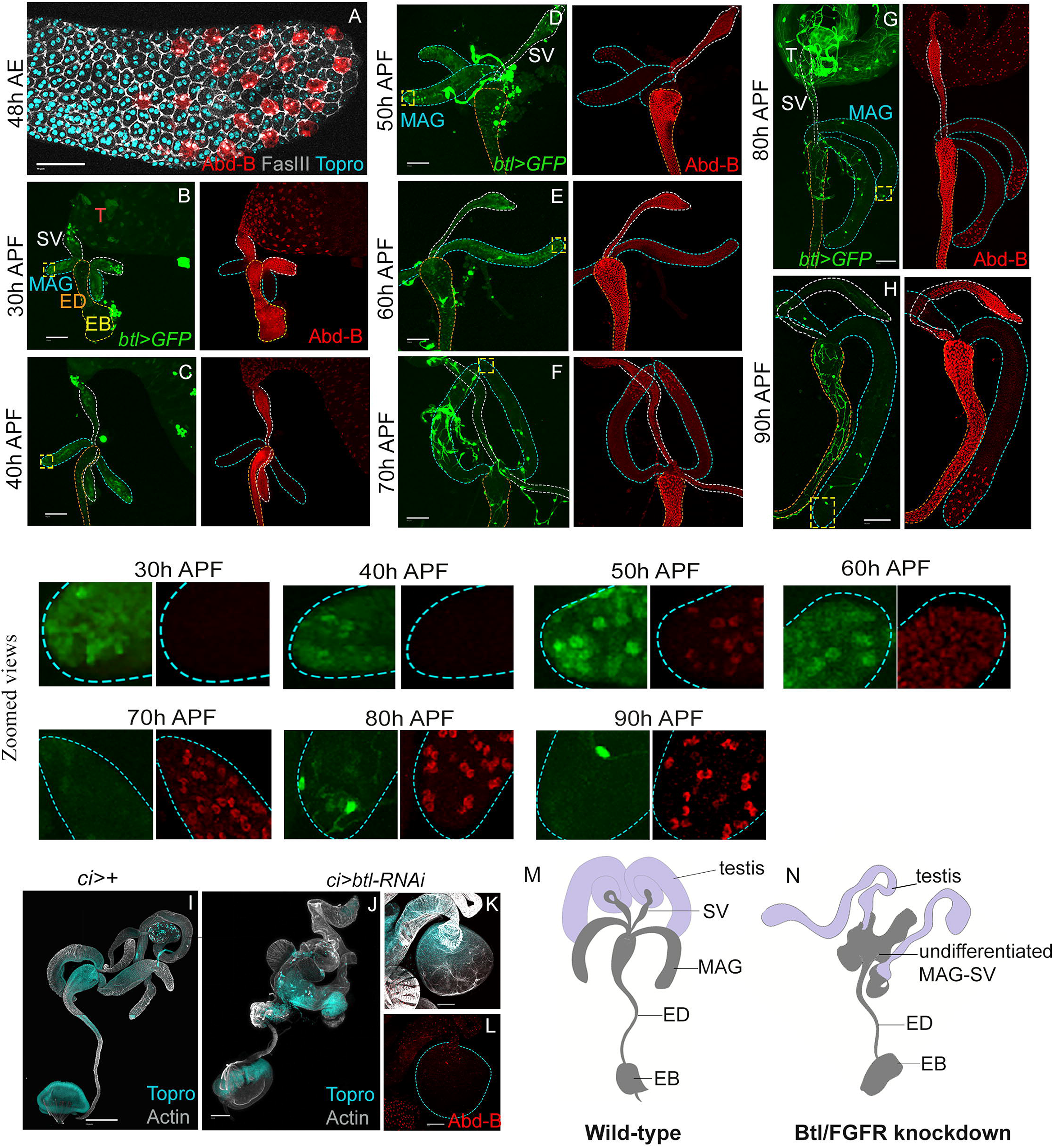
Spatio-temporal expressions of Btl/FGFR and Abd-B during pupal MAG morphogenesis. (A) Cell outlines of MAG epithelium (Fas III, grey) and cell nuclei (Topro, cyan) from a 2 day old adult after eclosion (AE), secondary cells are marked by Abd-B (Abd-B>GFP, red psuedocolor). (B-H) Expression pattern of Btl (*btl-Gal4>UAS-GFP*) and Abd-B (red) in the MAG-SV primordium derivates, MAG and SV, are shown from 30-90h APF. Distal tips in each image are shown at higher magnification in the bottom panel. Bright threads in D-H are the trachea. (I) Control *(ci-Gal4)* male reproductive system compared to (J) knockdown of *btl (citalic>btl-RNAi)* during MAG development marked by Actin (grey) and Topro (cyan). (K-L) shows a sample truncated MAG upon Btl knockdown, which displays loss of (L) Abd-B marked secondary cells in it. (M-N) Cartoon renditions of (M) control and (N) Btl knockdown are shown on the extreme right. Scale bar: 100μm Abbreviations and marking: testis (T), male accessory gland (MAG, blue broken line), seminal vesicle (SV, white dotted line), ejaculatory duct (ED, orange dotted line), ejaculatory bulb (EB, yellow dotted line).

These parallels between mammalian prostate and *Drosophila* MAG therefore hinted at similarities in their developmental ground plan. To further explore and elaborate this ground plan of the developing MAG, we examined Btl *(btl-Gal4>UAS-GFP)* and Abd-B during the entire course of pupal morphogenesis of MAG. We noticed that Btl/FGFR robustly marks the MAG bud early during its morphogenesis (30-50h APF, Figure 2B-D). Further, around 50h APF, Btl/FGFR displays robust expression in a group of distal cells (Figure 2D, see zoomed distal tip 50h APF) and, extinguishes, shortly ahead of the conclusion of pupal development (Figure 2E-H, 60-90h APF) suggesting its role during MAG bud growth and morphogenesis rather than in the adult gland.

The cell fate determinant posterior *Drosophila* Hox, Abd-B was not expressed in the growing MAG bud (30-50h APF, Figure 2B-D), unlike its expression in the developing SV throughout its budding and growth phase (Figure 2 B-H). Abd-B first appeared in the presumptive secondary cells of the MAG at 50h AFP (Figure 2D, see zoomed distal tip at 50h APF). Interestingly, Abd-B also initially expresses in the MAG main cells at 60h APF (Figure 2D, see zoomed distal tip at 60h APF) but persists only in the secondary cells through subsequent stages (Figure 2D-H, also see zoomed distal tips); thus its dynamic pattern of expression is marked by two characteristics: that is, its transient but ubiquitous expression in the entire gland, followed by its restricted expression in only the secondary cells.

Taken together, Btl/FGFR appears to be a primary toolkit for MAG-SV bud growth—a possibility that was further strengthened by the observation that knockdown (*ci-Gal4>btl-RNAi*) of Btl/FGFR truncated growth of the MAG (Figure 2J-K) with accompanying loss of its cell differentiation as revealed, for instance, by loss of development of the Abd-B-expressing secondary cells (Figure 2L). These features are reminiscent of fallouts of FGFR knockdown in prostate precursor cells in embryonic mouse, which completely abolishes growth of its anterior and ventral lobes and compromises growth of its dorso-lateral prostatic lobes as well as their cell fate acquisitions via. *Hoxd13* expression or other markers such as Bmp4 and TGF-β (Lin et al., 2007).

Btl/FGFR signaling and posterior Hox toolkit expression pattern in the highly branched and multi-lobular mammalian prostate (Huang et al., 2007) therefore appears to be a reiterative deployment of a simple tubular developmental ground plan of *Drosophila* MAG revealing deep homology of such accessory sexual organs from distant phylogeny.

### Hedgehog-Dpp signaling axis in MAG morphogenesis

In mouse prostate development, Shh signaling activity appears at budding, marks the nascent prostatic bud, declines as growth proceeds, and is seen in the gland of the adult gland at very low levels (Doles et al., 2006; Lamm et al., 2002). The ligand (Shh) is enriched at the distal prostatic tips of the epithelial bud, the receptor (Ptc), transcription factor (Gli) and downstream targets (e.g. Bmp4) are prominently expressed in the surrounding UGS mesenchyme (Lamm et al., 2001; Lamm et al., 2002; Podlasek et al., 1999; Pu et al., 2004). Developmental design reflected by Shh signaling with respect to prostate development, however, remains ambiguous owing in part to genetic redundancy: for instance, Gli1 and Gli3 may compensate for Gli2 loss in UGS tissues (Doles et al., 2006). It has, however, not been ascertained yet if MAG displays expression of the Hh/Shh signaling tool kit, reminiscent of that seen in prostate.

We thus examined the expression of Hh pathway members in the MAG bud; Ci (*ci-Gal4>UAS-GFP*), Ptc (*ptc-Gal4>UAS-GFP)* and Dpp *(dpp-Gal4>UAS-GFP*)—a Hh target, and mammalian Bmp counterpart—during the course of its pupal morphogenesis. While *ci>GFP* expression may not represent its actual activation—which is denoted by its cleaved form Ci-75 (Huangfu and Anderson, 2006; Price and Kalderon, 1999) —we consider it relevant for our purpose. We noted Ci expression around 20h APF: that is, during budding of MAG, (Figure 3A, see zoomed box, A’), which continued through its bud growth (Figure 3D, 30h APF) and cell differentiation stages (Figure 3G, 50h APF) and was finally extinguished in the MAG (Figure 3J, 90h APF). The spatio-temporal expressions of Hh receptor Ptc, too, marked the budding (20h APF, Figure 3B zoomed box), bud growth (30h APF, Figure 3E) and cell differentiation stages (50h APF, Figure 3H) albeit more prominently at the distal end compared to that of Ci. Moreover, unlike Ci that extinguishes by the end of pupal development, Ptc was retained in the secondary cells (90h APF, Figure 3K) suggesting its further roles in these cells in the adult gland. Hh target, Dpp, however, did not mark the nascent bud (20h APF, zoomed bud Figure 3C). It faintly appeared at the distal end during bud growth (30h APF, Figure 3F) and cell differentiation (50h APF, Figure 3I). Its robust expression was noted in the secondary cells by the completion of pupal MAG morphogenesis (90h APF, Figure 3L).

**Figure 3.**
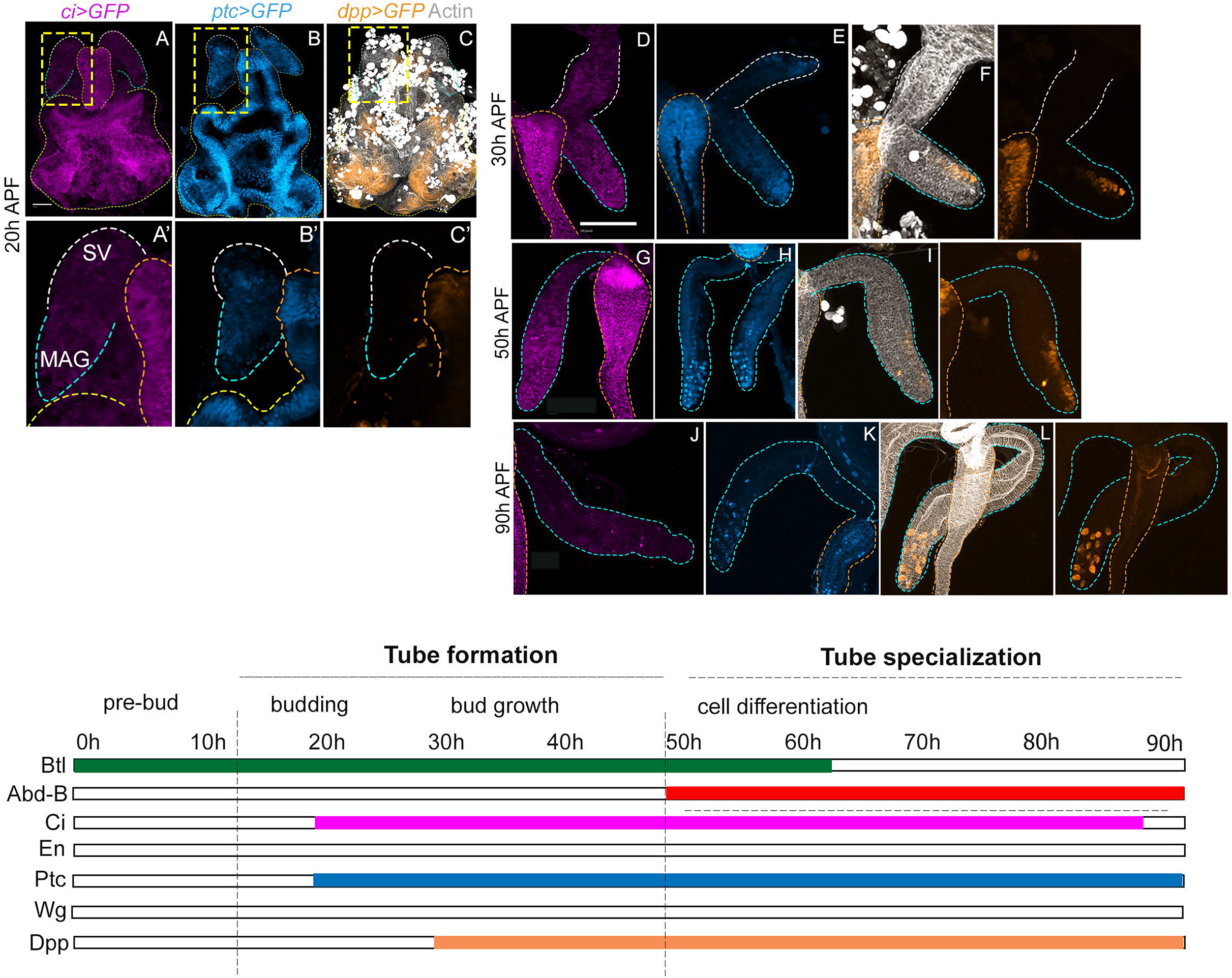
Hedgehog-Dpp signaling axis during MAG morphogenesis. (A-L) Expression of *ci* (A,D,G,J, purple), *ptc* (B,E,H,K, cyan blue) and *dpp* (C,F,I,L, orange) visualized by expressions of respective Gal4 driven UAS-GFP transgenes during budding (20h APF, boxed areas zoomed below), bud growth (30h APF) and cell differentiation (50 and 90h APF) stages. Diagram below summarizes the model of MAG developmental ground plan based on its toolkit expressions. Btl/FGFR is hallmark of the tube formation and Abd-B of tube specialization stage, while Hh-Dpp signaling toolkit is common to both stages. Scale bar: 100μm Marking scheme of the individual organs are as in preceding Figure 2.

These expression patterns during the course of MAG development therefore suggest Hh signaling requirement during bud growth stage and cell fate specification although Btl/FGFR may be the earliest crucial toolkit. While these inferences need critical functional evaluation, what appears most relevant to the current investigation is the fact that Hh signaling pattern in MAG development does appear reminiscent of that of prostate on the following general account: namely, its appearance in the nascent bud, followed by distal enrichment as the tube grows and, finally, decline in its expression upon formation of the tube.

## Conclusion

Co-options of common development ground plans via recruitment of shared toolkit genes mirror deep homology amongst organs of diverse phyla (Shubin et al., 1997; Shubin et al., 2009). Our study suggested that the developmental ground plan of MAG as a simplified tube also recurs during mammalian prostatic bud development, which are branched and turn into tubules via. canalization (Figure 4). These parallels reveal that in a blind watch-maker’s (Dawkins, 1986) analogy of evolution, common toolkits are co-opted across distant phylogenies, presumably *de novo*, which recreate a ground plan for development of organs. Reiterative deployment of this ground plan further results in modular designs of these organs, while deviations in these modules adds the diversities encountered across distantly related phylogenies (Wagner, 2007; Wagner and Altenberg, 1996). Binucleation and cuboidal-to-squamous transitions in the MAG epithelium (Taniguchi et al., 2014; Wilson et al., 2017) and columnar luminal cells, besides appearance of progenitor neuroendocrine cells in the mammalian prostate (Marker et al., 2003; Toivanen and Shen, 2017) represent individual phylogenetic peculiarities of these accessory sexual organs. Our study further suggests that—these evolutionary lineage specific peculiarities notwithstanding—MAG could also serve a model of a primitive prostate and therefore it could indeed be examined for modelling prostatic diseases, including cancers, for instance (Rambur et al., 2020).

**Figure 4.**
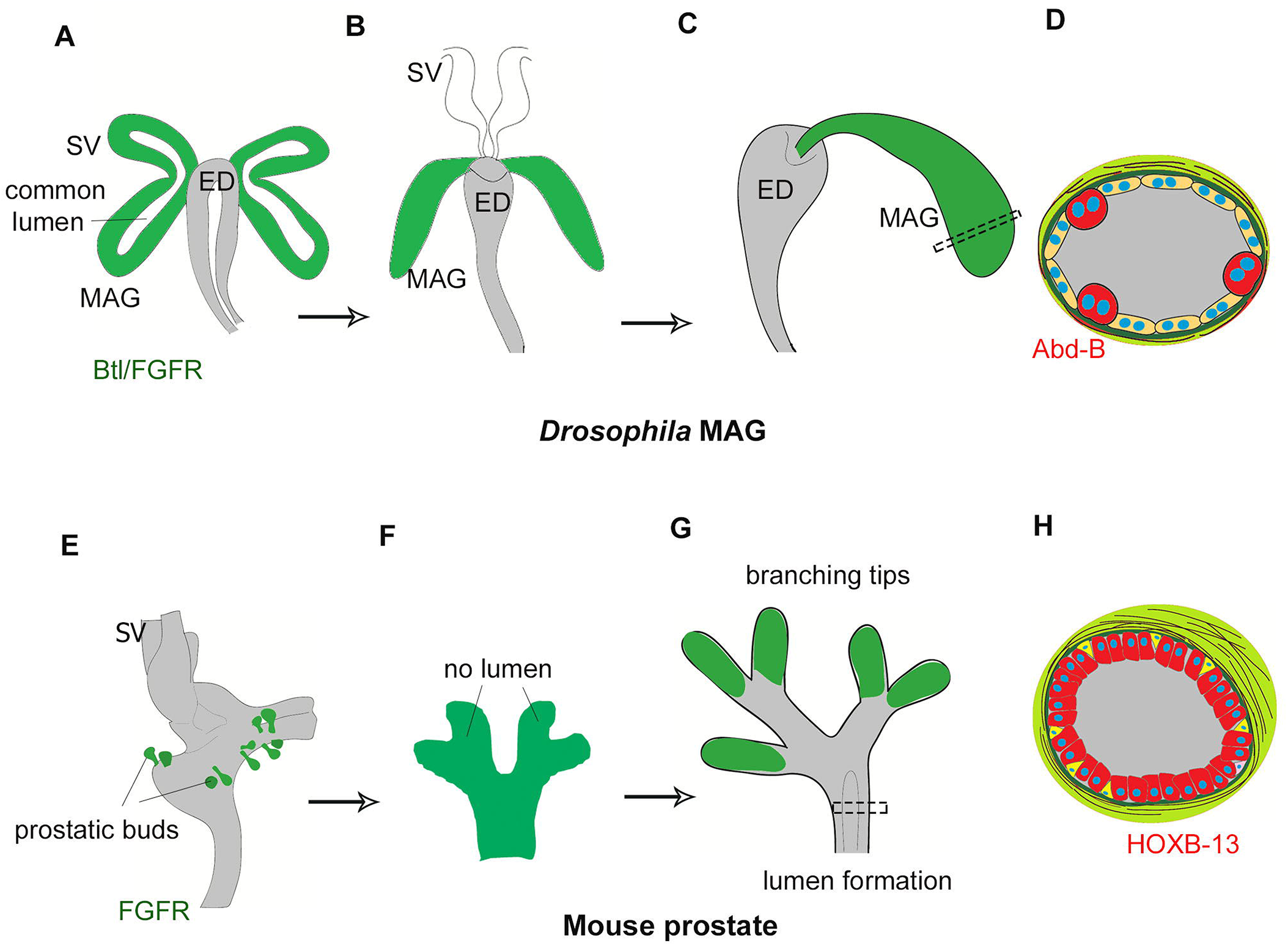
Common developmental ground plan of *Drosophila* MAG and mammalian prostate. (A-C) Cartoon representation of MAG development from Btl/FGFR primordial cells: (A) unitary MAG and SV buds with *a priori* lumen appear from the MAG-SV primordium. (B) MAG grows as a short tube. (C) MAG eventually connects caudally with the ejaculatory duct. (D) Illustration of MAG cross-section with two differentiated cell types, secondary cells are marked by Abd-B (E-G) Cartoon representation of mouse prostate development: (E) prostatic buds and SV are separate (F) individual primary prostatic bud does not have a lumen (G) branching with each individual tips marked by FGFR, while canalization precedes to form their lumen (H) Illustration of prostatic cross-section depicting two differentiated cell types, luminal cells are marked by Hoxb13.

## Material and Methods

### Collection and staging of pupal samples

Male pupae were identified by their prominent ovoid gonads. Males of the required genotype were selected by examining GFP expression under a fluorescent microscope. The 0h was decided as previously described (Bainbridge and Bownes, 1981). The maximum time variation may range ±30 minutes amongst individuals of a given stage. Collected 0h samples were allowed to grow at 25±1□ in a petri-dish with moist filter paper until the time of dissection. At least 5 individuals of a genotype were dissected for a given time point.

### Dissection of pupal MAG samples

Pupal dissection for 0-20h APF samples were done by excising the lower 1/3^rd^ of the pupae using dissection scissors in cold PBS. The genital disc was gently pushed out of the pupal cover using insulin syringes. For 30-70h pupal samples, first a cut was made at the abdomen-thorax junction. The reproductive structures within the abdomen were gently pushed out using insulin syringes. The tissue was cleared of abdominal fat and lipids. The pharate stage animals between 80-90h APF were taken out of the pupal case. MAG was isolated by gently removing the external genitalia using a pair of insulin syringes similar to adult dissection.

### Immunofluorescence staining and microscopy

Standard immunofluorescence protocol was followed. In brief, samples were fixed in 4% paraformaldehyde for 35 minutes, washed with PBS and incubated with the desired primary antibody overnight. The samples were washed, blocked in bovine serum albumin for an hour, before adding secondary antibody. Samples were counterstained with TO-PRO-3 (Invitrogen) and/or Phalloidin. Primary antibodies used: Mouse anti-En (1:50), Mouse anti-Abd-B (1:50), Mouse anti-FasIII (1:100); Phalloidin 555, 633 (1:100), TOPRO (1:500). Finally, the prepared samples were mounted in Vectshield or Prolong Gold for imaging. Images were acquired using a Leica confocal SP5 system and processed using LAS AF software.

### Key Resources Table

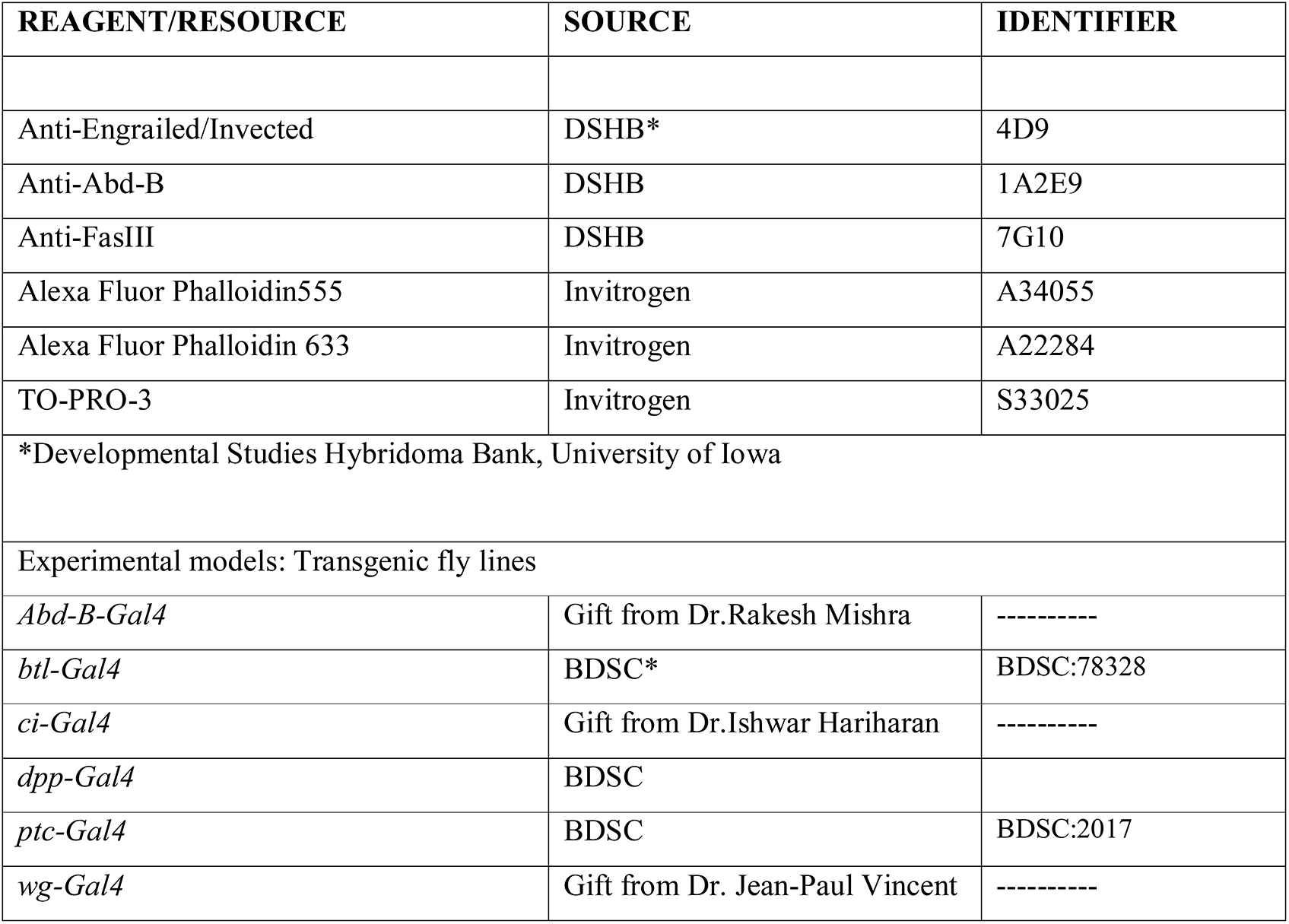
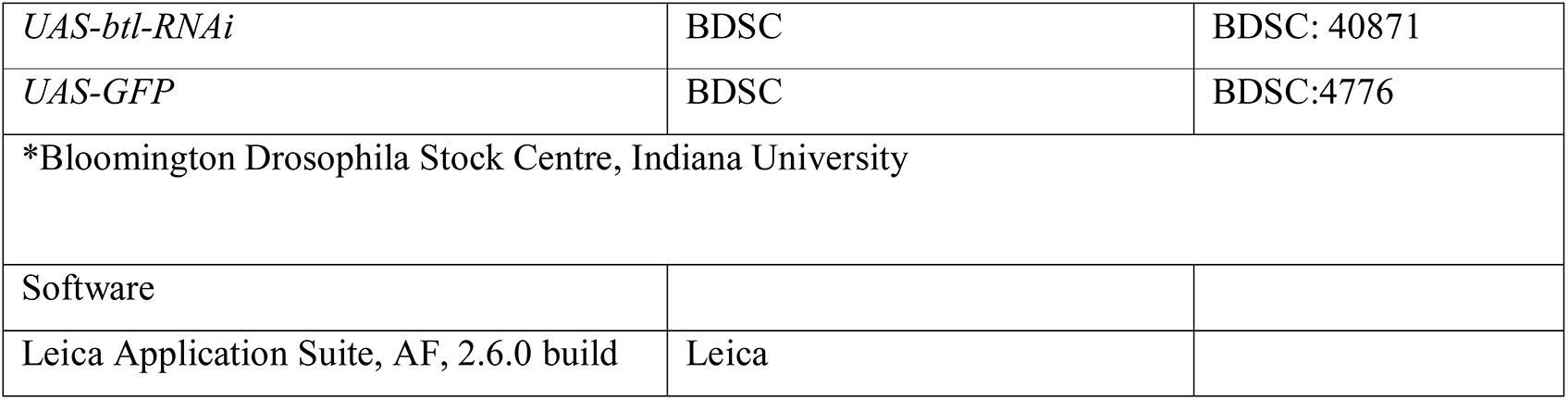

